# Effects of face masks on acoustic analysis and speech perception: Implications for peri-pandemic protocols

**DOI:** 10.1101/2020.10.06.327452

**Authors:** Michelle Magee, Courtney Lewis, Gustavo Noffs, Hannah Reece, Jess C. S. Chan, Charissa J. Zaga, Camille Paynter, Olga Birchall, Sandra Rojas Azocar, Angela Ediriweera, Marja W. Caverlé, Benjamin G. Schultz, Adam P. Vogel

## Abstract

Wearing face masks (alongside physical distancing) provides some protection against infection from COVID-19. Face masks can also change how we communicate and subsequently affect speech signal quality. Here we investigated how three face mask types (N95, surgical and cloth) affect acoustic analysis of speech and perceived intelligibility in healthy subjects. We compared speech produced with and without the different masks on acoustic measures of timing, frequency, perturbation and power spectral density. Speech clarity was also examined using a standardized intelligibility tool by blinded raters. Mask type impacted the power distribution in frequencies above 3kHz for both the N95 and surgical masks. Measures of timing and spectral tilt also differed across mask conditions. Cepstral and harmonics to noise ratios remained flat across mask type. No differences were observed across conditions for word or sentence intelligibility measures. Our data show that face masks change the speech signal, but some specific acoustic features remain largely unaffected (e.g., measures of voice quality) irrespective of mask type. Outcomes have bearing on how future speech studies are run when personal protective equipment is worn.

## A. INTRODUCTION

Face masks (alongside physical distancing) provide some protection against infection from Coronavirus disease (COVID-19) (Chu *et al*., 2020). Their use in public spaces and healthcare settings is either recommended or mandatory in many jurisdictions internationally. In the United States, the Center for Disease Control (CDC, 2020) recommends mask use to minimize droplet dispersion and aerosolization of the virus (Bahl *et al*., 2020). Clinical trials and healthcare settings continue to assess speech production, which generates respiratory droplets while unrestricted exposure increases the likelihood of disease contraction (Stadnytskyi *et al*., 2020). Risk of transmission increases through behaviors common in many speech assessment tasks including continuous and loud speech (Asadi *et al*., 2019). At the same time, acknowledgement of the necessity of personal protective equipment to minimize virus transmission has increased internationally (Asadi *et al*., 2019; Stadnytskyi *et al*., 2020; Zaga *et al*., 2020). Masks, however, alter the speech signal with downstream effects on intelligibility of a speaker. The use of personal protective equipment poses some unique challenges for speech assessment.

We evaluated the impact wearing a mask has on acoustic output and speech perception. We examined how different face mask types (surgical, cloth and N95), in combination with microphone location variations (headset vs. tabletop), affect speech recordings and intelligibility.

## B. METHODS

Four subjects, aged 29.0 ± 5.8 years, range 23-38; 2 males: 2 females, were included in the study. All speakers were English speaking with no dysphonia, cognitive or neurological impairments. One male and female had English as their second language.

### 1. Speech Acquisition

The speech battery was elicited by trained staff and consisted of sustaining an open vowel /aː/ for approximately six seconds reproduced ten times and reading a phonetically balanced text, the Grandfather Passage (Van Riper, 1963), reproduced five times. The speech battery was repeated under four conditions in a randomized order: 1) no mask; 2) standard surgical mask (regulated under 21 CFR 878.4040); 3) cloth mask (2-layered cotton); and 4) N95 mask (disposable mask made from electrostatic non-woven polypropylene fiber containing a filtration layer). Subjects were instructed to speak in a natural manner at a comfortable pitch and pace. Speech samples were recorded using two standardized methods: 1) Using a head-mounted cardioid condenser microphone (AKG520, Harman International, United States) positioned 2 inches from the corner of the subject’s mouth (minimum sensitivity of −43dB, near flat frequency response) and coupled with a QUAD-CAPTURE USB 2.0 Audio Interface (Roland Corporation, Shizuoka, Japan) connected to a laptop computer; and 2) Using a Blue Yeti (Blue Microphones, United States) tabletop microphone (sensitivity 4.5mV/Pa) connected to a laptop computer. The microphone was positioned 5 ft. from the subject to simulate physical distancing measures. Standardization of the recording environment was achieved by recording in the absence of traffic, electrical, appliance, or other background noise. All recordings were sampled at 44.1 kHz with 32-bit quantization.

### 2. Speech Intelligibility Testing

Speech intelligibility was evaluated using the Assessment of Intelligibility of Dysarthria Speech (ASSIDS) (Yorkston and Beukelman, 1984). For each condition subjects read aloud a randomized list of single words (one and two syllables in length) and sentences (5 to 28 syllables in length). Two blinded raters transcribed ASSIDS words and sentences, with the percentage of correct items calculated for each condition.

### 3. Audio processing and acoustic analysis

Audio files were screened for deviations and synchronized between microphones to ensure uniformity of length. Acoustic analysis of sustained vowel and reading tasks were performed using Praat software (Boersma, 2002). Two groups of speech features were analyzed, one to describe responsiveness to speech and silence, and another to determine agreement between measurements taken by different microphone conditions. The speech spectrum was used to describe the impact of mask type on the complex voice waveform. The interaction between intensity and frequency was characterized using the power spectral density (PSD, dB/kHz relative 2×10^−5^ Pa) in the longterm average spectrum on the reading task. PSD provides information on how “each frequency” contributes to the total sound power. Frequency bands were fixed at 1kHz. PSD was averaged across subjects for each mask condition and compared between masks not subjects.

Center-of-gravity (CoG, in Hz) was calculated from the power spectrum to inform frequency responsiveness of the conditions. CoG is the mean power-weighted frequency, i.e. the frequency that divides the power spectrum in equal halves above and below CoG. The intensity of background noise (floor) was determined as equal to the average intensity during the quietest three seconds of each files (i.e., in the absence of vocalization). Floor intensity was subtracted from the average intensity (during vocalization) for each task (vowel and reading) to determine the speech intensity prominence per mask condition. Features of interest included cepstral peak prominence smoothed (CPPS), harmonic-to-noise ratio (HNR), local jitter and shimmer for the sustained vowel, and average and standard deviation of pause length for the reading task.

Fundamental frequency was calculated through autocorrelation within a restricted range (70Hz - 250Hz for males, 100Hz - 300Hz for females) (Vogel et al., 2009). The analysis window was 43ms and 30ms respectively, and window shift fixed at 10ms. The maximum number of formants was set at 5 with a maximum of 5500Hz for formant detection. All other parameters were maintained at default software settings. The detection of silence-speech and speech-silence transitions was done using an energy threshold on the time domain (Rosen *et al*., 2010; Vogel *et al*., 2017). The threshold was set to 65% of the 95^th^ percentile, with minimum silence length set to 20ms and minimum speech length to 30ms.

### 4. Statistical analysis

To examine differences of each acoustic parameter under each mask condition (no mask, surgical, N95, and cloth), a linear mixed-effects model analysis using restricted maximum likelihood estimation was applied. Mask type was modeled as a fixed factor, and subject and order of mask as a random factor. Bonferroni corrected *post hoc* pairwise comparisons were conducted to determine differences in mask type (surgical, N95, and cloth) compared to no mask. To investigate power spectral density, the interaction effect between mask and frequency band was investigated. Where the interaction was significant, planned comparisons were made for each 1Khz frequency band to determine differences between masks types compared to no mask. SPSS was used for all statistical analyses (IBM SPSS Version 26.0).

## C. RESULTS

### 1. Speech intelligibility outcomes

Intelligibility varied between the speakers and across mask conditions. On average, intelligibility remained above 92% for all mask conditions, irrespective of single words (Figure 1a) or sentence tasks (Figure 1b). Single words (X=95.125±1.09) were perceived less accurately than words within sentences (97.25±0.645), (t=3.3128, *p*=0.0161). Average percentage correct scores were used for interpretation. Intelligibility for sentences for all conditions was between 97-98% accurate.

**FIGURE 1.**
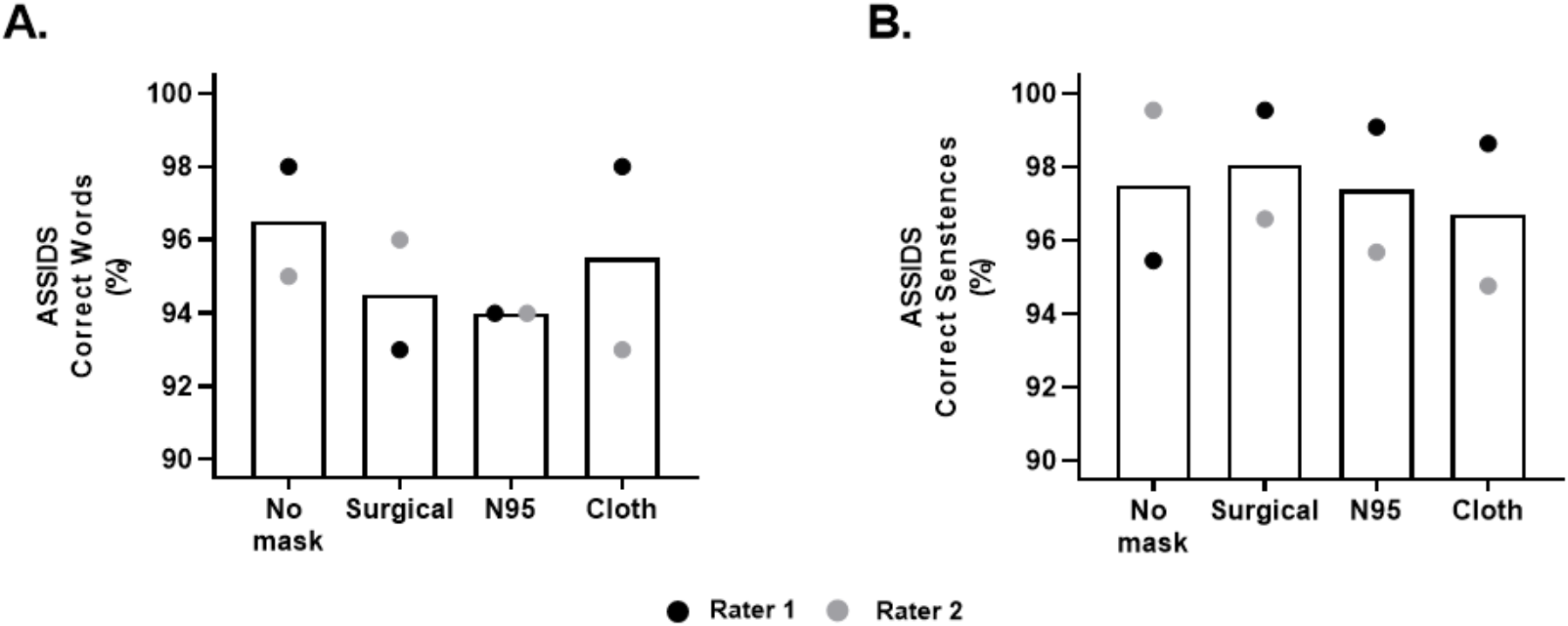
ASSIDS correct words and sentences each mask condition.

### 2. Power Spectral Density extracted from reading task under different mask conditions

Frequency bands were collapsed into 1kHz slices to explore differences in PSD between mask type. There was a Mask × 1kHz frequency band interaction effect (F_27,755_=2.50, *p*=0.006). Post hoc comparisons showed power (dB/Hz^2^) was significantly lower between 3-10 kHz for N95 mask and 5-10kHz for surgical and cloth masks when compared to no mask on recordings made using the head-mounted microphone (Figure 2a). No significant differences were observed between mask conditions on recordings made using the tabletop microphone (F_27,757_=1.41, *p*=0.082; Figure 2b).

**FIGURE 2.**
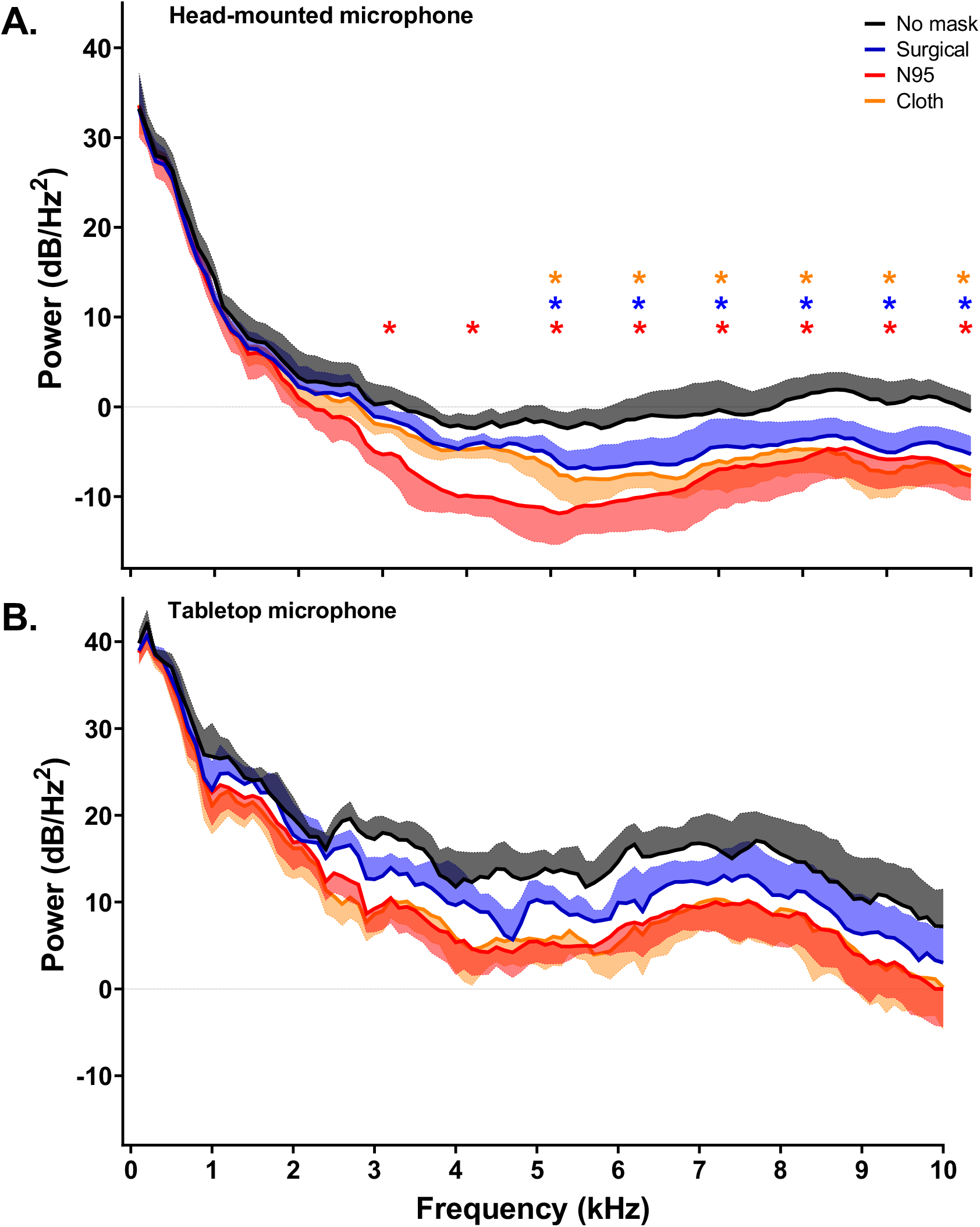
Power spectral density extracted from reading task under different mask conditions. Mean power spectra density displayed between 1-10kHz based on mask type. Shaded areas represent the standard error of mean. *p≤0.05 no mask vs mask type at each frequency bin. Red stars denote significant differences between no mask and N95, blue stars denote significant differences between no mask and surgical masks while orange stars denote significant differences between no mask and N95.

### 3. Acoustic parameters extracted from sustained vowel and reading tasks

For recordings produced with the head-mounted microphone, there was a significant effect of masks for mean pause length (F_3,8.97_=3.88, *p*=0.05), percentage of pauses (F_3,8.40_=7.36, *p*=0.01) and spectral tilt (F_3,8.98_=13.62, *p*=0.001) extracted from the reading task. Post hoc comparisons showed that recordings produced with the N95 mask increased percentage of pauses (*p*=0.023) (Table 1). Spectral tilt was lower in recordings produced with the surgical (*p*=0.016) and N95 masks (*p*=0.001). For recordings produced with the tabletop microphone, there was a significant effect of mask type for percentage of pauses (F_3,7.87_=8.17, *p*=0.008), and spectral tilt (F_3,8.39_=15.43, *p*=0.001) (Table 1). Post hoc comparisons revealed that the N95 and cloth masks yielded higher percentage of pauses (N95 *p*=0.022; Cloth *p*=0.029) no mask. As with the head-mounted microphone, recordings produced with the tabletop microphone yielded lower spectral tilt values with both the surgical (*p*=0.006) and N95 masks (*p*=0.002). No significant differences were observed in acoustic parameters extracted from the sustained vowel recorded using either the headmounted or tabletop microphone.

**TABLE 1.** Acoustic parameters extracted from the reading task recordings produced by the head-mounted and tabletop microphones under different mask types.

|  | No Mask | Surgical | N95 | Cloth | *F* | Mean Diff (95% CI) |  |  |
| --- | --- | --- | --- | --- | --- | --- | --- | --- |
|  |  |  |  |  |  | No Mask<br>vs. Surgical | No Mask<br>vs. N95 | No Mask<br>vs. Cloth |
| **Head-mounted microphone** |  |  |  |  |  |  |  |  |
| Mean pause length (seconds) | 0.24 ± 0.07 | 0.24 ± 0.08 | 0.28 ± 0.10 | 0.26 ± 0.10 | **3.88\*** | 0.008<br>(-0.053, 0.036) | 0.032<br>(-0.012, -0.077) | 0.019<br>(-0.025, 0.063) |
| Variability of pause length | 0.36 ± 0.09 | 0.38 ± 0.14 | 0.44 ± 0.16 | 0.43 ± 0.17 | 3.14 |  |  |  |
| Percent of pauses (%) | 30.3 ± 3.88 | 31.74 ± 2.56 | 35.42 ± 3.08 | 34.94 ± 2.76 | **7.36\*\*** | 1.00<br>(-3.23, 5.22) | **4.91\***<br>**(0.66, 9.17)** | -4.25<br>(-0.02, 8.52) |
| Spectral tilt (dB) | -21.4 ± 3.32 | -16.73 ± 1.84 | -14.5 ± 3.06 | -18.86 ± 3.65 | **13.62\*\*\*** | **4.65\***<br>**(0.83, 8.47)** | **6.92\*\*\***<br>**(3.10, 10.74)** | 2.49<br>(-1.33, 6.31) |
| Mean Intensity (dB) | 63.61 ± 3.04 | 63.04 ± 3.35 | 63.27 ± 3.75 | 62.05 ± 3.05 | 1.50 |  |  |  |
| Intensity prominence | 42.86 ± 2.03 | 40.66 ± 3.07 | 41.68 ± 2.18 | 40.01 ± 2.73 | 2.68 |  |  |  |
| p95 Intensity | 64.37 ± 3.01 | 63.83 ± 3.37 | 64.05 ± 3.72 | 62.97 ± 2.97 | 1.19 |  |  |  |
| CPPS | 19.40 ± 2.89 | 20.58 ± 1.76 | 20.8 ± 2.57 | 20.31 ± 2.75 | 2.21 |  |  |  |
| HNR | 24.68 ± 3.45 | 25.48 ± 3.23 | 25.84 ± 5.09 | 26.56 ± 3.78 | 1.41 |  |  |  |
| f0 MEAN (Hz) | 155.42 ± 63.82 | 155.09 ± 66.08 | 154.1 ± 63.90 | 162.25 ± 60.69 | 0.92 |  |  |  |
| f0 CoV (%) | 0.76 ± 0.07 | 0.72 ± 0.08 | 0.63 ± 0.08 | 0.72 ± 0.09 | 2.60 |  |  |  |
| jitter (%) | 0.31 ± 0.07 | 0.36 ± 0.09 | 0.31 ± 0.07 | 0.34 ± 0.09 | 1.45 |  |  |  |
| shimmer (%) | 1.51 ± 0.23 | 1.55 ± 0.16 | 1.64 ± 0.5 | 1.51 ± 0.24 | 0.49 |  |  |  |
| **Tabletop microphone** |  |  |  |  |  |  |  |  |
| Mean pause length | 0.38 ± 0.16 | 0.40 ± 0.17 | 0.41 ± 0.18 | 0.42 ± 0.20 | 0.80 |  |  |  |
| Variability of pause length | 0.41 ± 0.13 | 0.46 ± 0.17 | 0.50 ± 0.17 | 0.50 ± 0.19 | 3.29 |  |  |  |
| Percent of pauses (%) | 25.37 ± 4.84 | 26.25 ± 4.50 | 29.04 ± 4.47 | 28.91 ± 5.56 | **8.17\*\*** | 0.80<br>(-2.21, 3.81) | **3.54\***<br>**(0.50, 6.57)** | **3.39\***<br>**(0.34, 6.44)** |
| Spectral tilt (dB) | -30.82 ± 1.43 | -24.78 ± 1.82 | -23.59 ± 4.09 | -29.32 ± 4.96 | **15.43\*\*\*** | **6.59\*\***<br>**(2.03, 11.15)** | **7.65\*\***<br>**(3.09, 12.21)** | 1.80<br>(-2.76, 6.35) |
| Mean Intensity (dB) | 71.54 ± 3.89 | 71.73 ± 4.34 | 71.85 ± 4.31 | 72.26 ± 2.78 | 0.12 |  |  |  |
| Intensity prominence | 37.09 ± 3.91 | 36.67 ± 4.35 | 36.94 ± 4.5 | 37.57 ± 3.12 | 0.22 |  |  |  |
| p95 Intensity | 72.66 ± 3.76 | 72.87 ± 4.3 | 72.95 ± 4.37 | 73.52 ± 2.84 | 0.19 |  |  |  |
| CPPS | 19.52 ± 2.74 | 19.16 ± 1.87 | 19.99 ± 2.19 | 19.34 ± 2.1 | 0.52 |  |  |  |
| HNR | 20.30 ± 3.66 | 19.11 ± 3.25 | 21.88 ± 3.77 | 21.37 ± 2.16 | 1.19 |  |  |  |
| f0 MEAN (Hz) | 155.80 ± 63.25 | 155.4 ± 64.64 | 156.4 ± 61.32 | 169.77 ± 45.03 | 0.92 |  |  |  |
| f0 CoV (%) | 0.71 ± 0.09 | 0.77 ± 0.08 | 0.65 ± 0.08 | 0.65 ± 0.06 | 2.41 |  |  |  |
| jitter (%) | 0.32 ± 0.06 | 0.36 ± 0.11 | 0.31 ± 0.08 | 0.32 ± 0.06 | 0.98 |  |  |  |

\* $p \leq 0.05$ , \*\* $p \leq 0.01$ , \*\*\* $p \leq 0.001$ , ± represent one standard deviation

## D. DISCUSSION

The type of mask affected the speech signal. We observed significant differences in acoustic power distribution across relevant frequency bands for speech in all three mask conditions compared to no mask. The differences were not observed in frequencies below 3kHz. Differences in signal for higher frequencies led to altered acoustic outcomes including spectral tilt. The masks however did not significantly influence listener-perceived intelligibility or acoustic measures of perturbation (e.g., NHR, CPPS). Measures of speech rate were lower for N95 and surgical masks, possibly as speakers compensate when wearing masks to improve intelligibility. It is also possible that speech timing differences were related to how speech boundaries are identified in the analysis scripts (i.e., our timing analysis relied on identification of phoneme/word boundaries via intensity thresholds).

Intelligibility scores varied between raters and between mask condition. Intelligibility remained above 92% for words and sentences. Anecdotally, it can be difficult to understand people when they wear a mask (Goldin et al., 2020). Our small dataset suggests mask type does not systematically impact intelligibility in controlled environments. Our recordings were made with high-quality microphones in quiet environments. Raters listened to samples in ideal listening conditions away from distractions and background noise but without visual aid (lips and jaw movement) for all mask conditions. In loud environments, communication can be challenging with multiple distractors, background noise, and a lower signal-to-noise ratios (SNR). Noise in ecological situations may further decrease speech intelligibility, when complementary visual cues blocked by use of face masks play a role in communication.

It is clear that face masks change the acoustic speech signal, but some specific perceptual features remain largely unaffected (e.g., acoustic measures of voice quality) irrespective of mask type. These results have implications for clinical assessments and speech research where PPE is required. It is easy to assume that subjects in a speech study will simply remove PPE during assessments; however, subjects and researchers may be reluctant to do so if it leads to potential exposure to airborne viruses. In longitudinal studies with data collection before, during, and after pandemics requiring PPE, researchers should consider how to mitigate against changes to protocols that affect speech (see Figure 3) (Redenlab, 2020).

**FIGURE 3.**
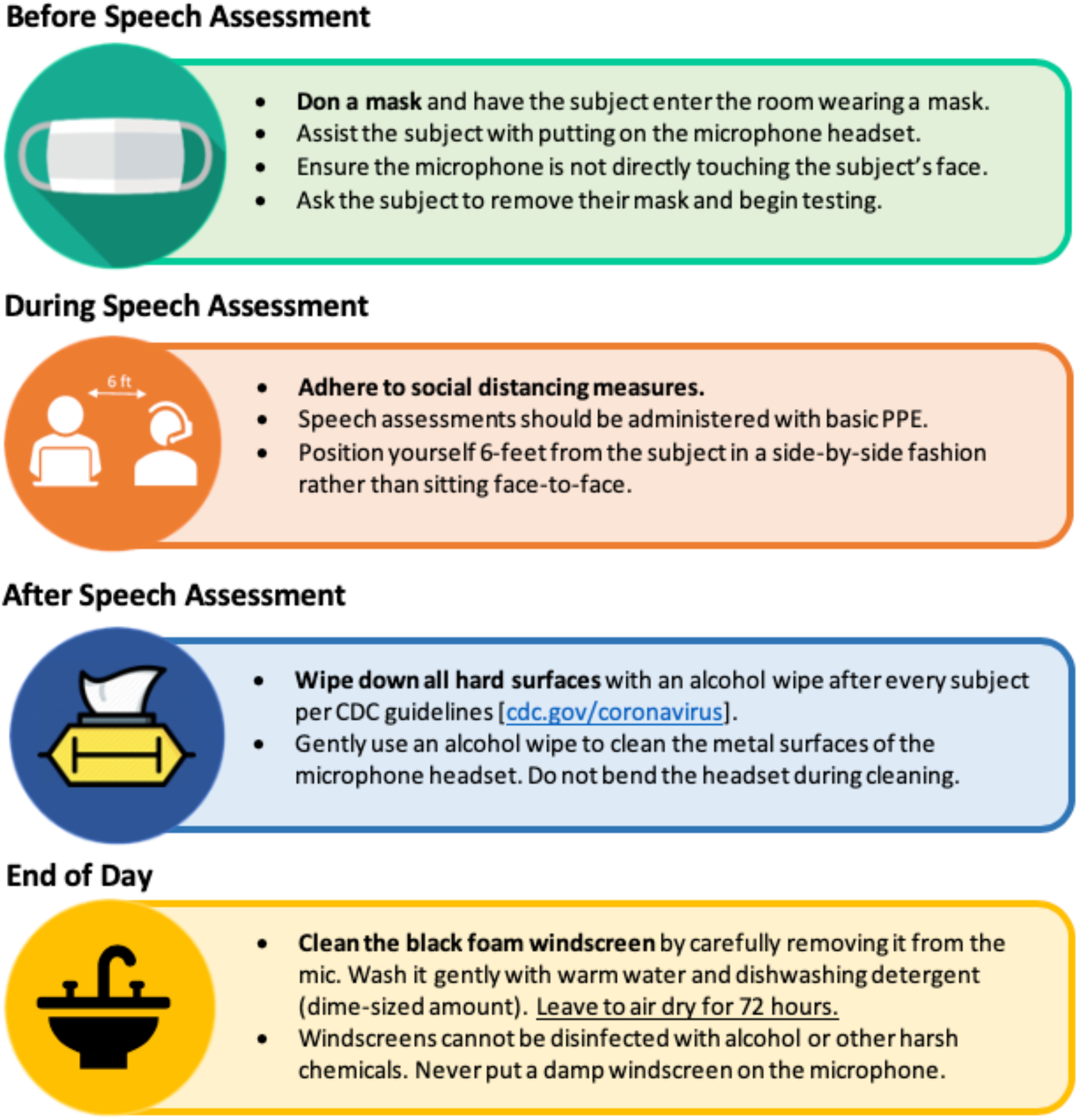
Guidance on minimizing risk to patients and staff during speech recordings (reproduced with permission from Redenlab Inc, https://redenlab.com/clinical-resources) To reduce risk, it is recommended assessors wear masks throughout assessments, the microphone’s metal surfaces are sanitized between subjects, and all windscreens are washed at the end of each use.

## E. ACKNOWLEGEMENTS

This work received institutional support from The University of Melbourne, Australia. APV holds a National Health and Medical Research Council (Australia) Fellowship (#10135683).

CL, GN, OB and MC are supported by Australian Postgraduate Research Scholarships. CP is funded by a joint National Health and Medical Research Council (Australia)/Motor Neuron Disease Research Australia postgraduate scholarship (#1133541)

***\*Disclaimer**: Please be advised that nothing completely eliminates bacteria or viruses and the guidelines contained in this document are measures attempting to limit the spread of a virus. Further, these guidelines do not supersede medical practitioner recommendations or the COVID-19 safety policies implemented by your business or institution. It is your responsibility to follow the recommendations and*

## Notes

https://redenlab.com/clinical-resources

### Competing Interest Statement

APV, CL and MM work for Redenlab, a speech analytics company.

